# HP1α-driven Phase Separation and Repair Pathway Choice in Response to Heterochromatin Damage

**DOI:** 10.1101/2024.09.16.613371

**Authors:** Darshika Bohra, Aprotim Mazumder

## Abstract

Double-strand breaks (DSBs) pose significant threat to genomic stability and need immediate attention from DNA Damage Response (DDR) machinery involved in Homologous Recombination (HR) or Non-homologous end joining (NHEJ). DDR in heterochromatin is challenging owing to the distinct chromatin organization. Heterochromatin Protein 1 (HP1) isoforms that contribute significantly to the organization of heterochromatin, have been shown to be involved in DDR. Mammalian HP1 has three isoforms, HP1α, HP1β, and HP1γ, which possess significant homology and yet have distinct functions. HP1α is the only isoform known to undergo liquid-liquid phase separation. We show that the minute-scale dynamics of HP1α and HP1β differ dramatically and they promote differential recruitment of HR vs. NHEJ factors at the sites of laser-induced clustered DSBs. Perturbing HP1α phase-separation abrogates both the recruitment of HR factors and readouts of HR. Our study provides a link between phase-separation and DDR-centric roles of HP1α and hints at spatial partitioning of repair pathways in response to damage in heterochromatin.

## Introduction

DNA-related processes such as replication, transcription, and repair occur in the context of chromatin (Dinant et al., 2008; Groth et al., 2007; B. Li et al., 2007). One of the main structural and functional distinctions of chromatin is the loosely packed euchromatin and compact heterochromatin (Hennig, 1999; Murakami, 2013). Heterochromatin is usually characterized by silencing histone methylation marks, which ensure compaction and inhibit transcription (Grewal & Jia, 2007). The main methylation mark is H3K9 trimethylation (H3K9me3), which is recognized by the major architectural protein of heterochromatin, the heterochromatin protein 1 (HP1) (Bannister et al., 2001; Grewal & Jia, 2007; Lachner et al., 2001). There are three isoforms of HP1 in mammals, HP1α, HP1β, and HP1γ (Canzio et al., 2014; Jones et al., 2000). The three isoforms have high homology in sections but also have non-redundant functions and localization (Canzio et al., 2014). HP1α and HP1β localize to heterochromatin, while HP1γ localizes to both euchromatin and heterochromatin (Jones et al., 2000). HP1α and HP1β have significant sequence homology in the chromodomain and chromoshadow domain but exhibit differences in the sequence, specifically in the N-terminal region of the protein, which brings about unique functions and modulates affinity to the H3K9me3 mark that these proteins bind (Canzio et al., 2014; Hiragami-Hamada et al., 2011). A major functional difference between the two isoforms has recently been discovered, i.e. the ability of HP1α to undergo the process of liquid-liquid phase separation (LLPS), which is thought to be important for heterochromatin formation and maintenance (Keenen et al., 2021; Larson et al., 2017; Strom et al., 2017). Independent studies have described the involvement of HP1 proteins in processes of DNA damage repair. In one study, HP1 proteins were found to have an inhibitory role in the repair of heterochromatic double-strand breaks (DSBs), and the knockdown of HP1 helped alleviate the inhibition and aid the repair process without the need for the master kinase ATM (Goodarzi et al., 2008). On the other hand, in a different study, the release of HP1β from the site of laser-induced damage occurred within seconds, and such HP1β mobility was necessary to induce the DDR signalling, in the absence of which the damage persisted (Ayoub et al., 2008). Additionally, it has been shown that all HP1 isoforms get recruited to the site of damage, and this was independent of the H3K9me3 marks, highlighting that the binding to damaged chromatin was independent of the canonical heterochromatin recruitment of HP1 isoforms (Luijsterburg et al., 2009). HP1α has also been shown to mediate DSB repair through HR (Baldeyron et al., 2011; Soria & Almouzni, 2013). Despite this the exact roles of different HP1 isoforms in DSB repair remain unclear. Moreover, the functions specifically in the context of DSBs in heterochromatin have not been explored as heterochromatin organization may offer specific challenges to the repair machinery (Cann & Dellaire, 2011; Fortuny et al., 2021; Janssen et al., 2018). It has been shown that DSB repair in heterochromatin requires specific and unique mechanisms owing to its architecture (Amaral et al., 2017). Specifically, DSBs in *Drosophila* heterochromatin migrated to the periphery of the heterochromatin domain before the recruitment of RAD51 and the onset of repair via HR pathway (Chiolo et al., 2011). Additionally, the DSBs were shown to further migrate to the nuclear periphery to complete the HR repair pathway, which was mediated by nuclear actin and myosin (Caridi et al., 2018). These special machinery exists to help relocalize the DSBs from the repetitive heterochromatin domain to prevent ectopic recombination events, failing which the genome stability is compromised (Caridi et al., 2018; Chiolo et al., 2011; Janssen et al., 2018). These studies suggest that the repetitive nature of heterochromatin sequence warrants distinctive dynamics to prevent illegitimate recombination and lethal chromosome rearrangements in the event of damage.

Therefore, it is important to explore the different ways in which heterochromatin DSB repair can occur without the risk of ectopic recombination events. The role of the recently discovered property of LLPS of HP1α in mediating repair also remains unexplored. In this study, we explore the differential roles of HP1α and HP1β in the repair of heterochromatic DSBs. Additionally, we investigate whether the phase separation of HP1α plays a role in such repair. Our data suggests that the two HP1 isoforms display significantly different localization dynamics in response to clustered DSBs in heterochromatin. Furthermore, they affect the differential recruitment of HR and NHEJ factors at the site of damage, indicating a critical role for HP1 isoforms in repair pathway choice and spatial partitioning of these processes. We also show that HP1α-mediated phase separation is critical for such spatial partitioning of heterochromatin DSB repair and for the timely resolution of the heterochromatin DSBs in a positionally stable manner.

## Results

### HP1 isoforms have differential dynamics at the site of laser-induced damage

The dynamics of HP1 isoforms have been studied using Fluorescence Recovery After Photobleaching (FRAP) (Cheutin et al., 2003; Schmiedeberg et al., 2004), even in the context of DNA damage (Ayoub et al., 2008; Zarebski et al., 2009). Most of these studies have focused on faster dynamics over several seconds, but given the recently described property of HP1α of undergoing phase separation (Keenen et al., 2021; Larson et al., 2017; Strom et al., 2017), and the general importance of LLPS in DDR (Altmeyer et al., 2015; Kilic et al., 2019; Wang et al., 2023) we wondered if the longer-term dynamics of HP1 isoforms may be different at sites of clustered DSBs in the heterochromatin (HC) corresponding to the setting up of a phase-separated domain conducive to repair. To address this, we expressed GFP-tagged HP1α or HP1β in U2OS cells. Routine FRAP involves using a laser (usually the wavelength that causes maximal excitation of the fluorophore the protein is tagged with) to photobleach a region of interest, which results in a loss of fluorescence signal. Bleaching is followed by tracking the fluorescence recovery over time to quantify the protein dynamics (Axelrod et al., 1976; Houtsmuller, 2005; Ishikawa-Ankerhold et al., 2012). We used a modified FRAP protocol where a 405 nm laser was used for photobleaching and inducing double-strand breaks in Hoechst-sensitized cells, as described below (also see Materials and Methods). Heterochromatin nodes can be identified by Hoechst staining and HP1 localization. To evaluate HP1 dynamics in the context of damage, we combined the process of FRAP with DSB induction using a 405 nm laser and Hoechst sensitization (Fig. 1A). This involves treating the cells with Hoechst for 10 minutes, which allows the induction of DSBs when the cells are exposed to a 405 nm laser, following and modified from earlier work (Kruhlak et al., 2006; Rogakou et al., 1999). We chose heterochromatin nodes from the GFP channel intensity and irradiated the region with the 405 nm laser using a point ROI (see Materials and Methods). Cells that are not treated with Hoechst should harbor no DSBs and only undergo photobleaching, thus serving as controls. In contrast, in cells that are treated with Hoechst, irradiation would cause photobleaching along with the induction of DSBs (Supp. Fig. 1A). The recovery of fluorescence was tracked over 10 minutes to get insight into the long-term dynamics of HP1 as opposed to the short-term dynamics that had been observed earlier (Ayoub et al., 2008; Cheutin et al., 2003). GFP-tagged HP1α and HP1β showed remarkably different dynamics after DSB induction (while they were quite comparable in control non-Hoechst cells). GFP-HP1α recovered within minutes, though the mobile fraction reduced upon damage; GFP-HP1β, on the other hand, recovered a few seconds after irradiation only to disperse precipitously soon after (Fig. 1A, B). It did not show a recovery to baseline levels over the next 10 minutes. The observed dynamics are a combination of both the protein dynamics and large-scale chromatin movement away from the damage site, as described before (Ayoub et al., 2008; Burgess et al., 2014). Thus, overall, the dynamics of GFP-tagged HP1 isoforms upon laser-induced clustered DSBs evidence a normal enrichment of HP1α at the site of damage and depletion of HP1β. Similar dynamics for GFP-tagged HP1α and HP1β were also observed in HeLa cells (Supp. Fig. 1B). To verify that this is not an artefact of overexpression of GFP-tagged proteins, we performed laser irradiation followed by immunofluorescence for endogenous HP1α and HP1β in cells transfected with PARP1-chromobody (tagRFP). PARP1-chromobody helps visualize endogenous PARP1 and thus allows us to monitor the site of damage in real-time without artefacts associated with the overexpression of PARP1. In these cells, HP1α showed accumulation at the site of damage, while HP1β did not show any enrichment in the same cells and was, in fact, partially depleted (Fig. 1C). HP1β seemed to accumulate at the periphery of the damaged foci, marked by the enrichment of PARP1 (Fig. 1C). This was quantified by taking a ratio of the mean intensity at the site of damage to the intensity in the nucleus. Such quantification clearly shows that HP1α shows preferential enrichment at the site of damage while HP1β is depleted (ratio less than 1). It accumulates at the periphery of the site of damage in some cells while remaining dispersed in other cells (Fig. 1C, E). This is captured both in the intensity line profile at the site of damage (Fig. 1D) and in the quantification across several cells (Fig. 1E). Poly-ADP-ribosylation has been known to drive phase separation at sites of clustered DSBs and play a critical role in the recruitment of repair factors ^29^. PARP gets recruited early to the site of damage, and thus, it is an ideal candidate for regulating HP1 proteins. We treated cells with the PARP inhibitor (PARPi), AZD2461, and performed laser irradiation followed by immunofluorescence for HP1α. We observed a multifold downregulation in PAR enrichment, suggesting that PARP inhibition occurred efficiently, but there was no significant difference in HP1α enrichment between the control and PARP-inhibitor treatments suggesting that PARP does not have a significant role to play in regulating HP1 dynamics in the event of damage (Fig. 1F). Thus, our data suggests that HP1 isoforms show distinct dynamics at clustered DSBs in heterochromatin, with HP1α showing enrichment while HP1β remains depleted.

**Fig. 1.**
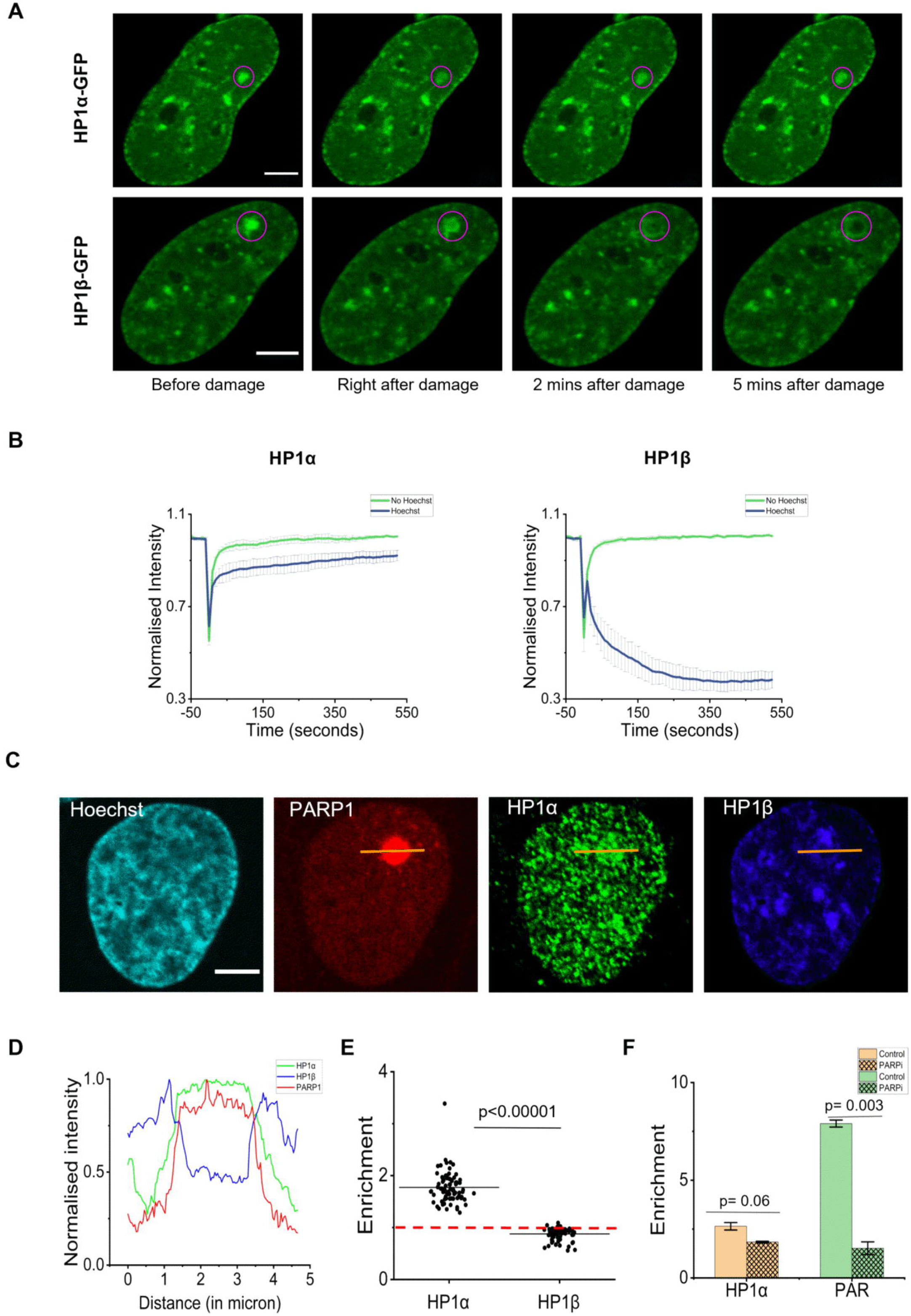
HP1 isoforms have differential dynamics at the site of damage. (A) Representative images showing dynamics of GFP-tagged HP1α and HP1β at different time points of a FRAP experiment with Hoechst sensitization. The magenta circle represents the heterochromatin node that was bleached/damaged with a 405 nm laser. The scale bar is 5μm. (B) FRAP curves showing the dynamics of GFP-tagged HP1α and HP1β in undamaged control (No Hoechst, green) and damaged cells (Hoechst, blue). The curve depicts the mean and standard error of mean (SEM) from three experiments (N=3, n>15 cells, each condition). (C) Representative images from immunofluorescence experiment showing endogenous HP1α (green) and HP1β (blue) staining at the site of damage marked by PARP1(red). The scale bar is 5μm. The yellow line represents the line across which intensity profile in (D) is plotted. (D) The intensity of PARP1, HP1α, and HP1β along the yellow line shown in 1C is plotted. The site of damage has higher intensities of PARP1 and HP1α while the intensity of HP1β is low. (E) The enrichment of HP1α and HP1β at the site of damage quantified from the image in 1C. is shown. The distribution of cells combined from three immunofluorescence experiments is shown (N=3, n>50 cell, each condition). The line depicts the mean. The p-value is calculated using the Kolmogorov-Smirnov test. Enrichment is defined as the ratio of intensity at the site of damage to the intensity in the whole nucleus. (F) Bar graph showing enrichment of HP1α (orange) and PAR (green) in control cells (undashed) and AZD2461 treated cells (dashed) show no significant difference. Means and standard deviations (SD) from two repeats are shown (N=2, n>60 cells, each condition). The p-value is calculated using the student’s t-test. Enrichment is defined as the mean of the ratios of intensity at the site of damage to the intensity in the whole nucleus.

In the dynamics experiments above, the behaviour of a heterochromatin node appears to be different depending on whether HP1α is overexpressed or HP1β. Strikingly, we also observed that cells that are overexpressing HP1α showed reduced staining for HP1β and vice versa (Supp. Fig. 1C). The quantification shows a clear downregulation of one isoform on the overexpression of another isoform (Supp. Fig. 1C). Since we wanted to tease apart the roles of the two isoforms in DDR, we used this feature to our advantage in subsequent experiments to understand how HP1α or HP1β may influence DDR.

### HP1 dynamics can directly influence repair pathway choice at the DSBs in heterochromatin

HP1α enrichment is important for promoting HR (Baldeyron et al., 2011), and HP1β depletion has been implicated in γH2AX induction (Ayoub et al., 2008); hence, we wondered if the differential localization of HP1α versus HP1β at sites of laser-induced DSBs may be correlated to differential DSB repair pathway choice at heterochromatin sites. We first investigated the major DDR protein 53BP1, a pro-NHEJ factor (Lei et al., 2022). We performed heterochromatin-specific irradiation in cells overexpressing HP1α or HP1β, followed by immunofluorescence for 53BP1. 53BP1 showed distinct localizations around the HP1α or HP1β foci. The localization was more homogeneous in cells overexpressing HP1β, while it seemed to enrich in a ring-like arrangement around the HP1α foci with a void where HP1α intensity was the highest (Fig. 2A). We quantified this by taking a ratio of the intensity at the site of damage to the intensity in an annulus right around the site of damage (Supp. Fig. 2A). This indicated that 53BP1 localizes directly at the site of damage in cells overexpressing HP1β but remains peripherally enriched around HP1α foci (Fig. 2A). The localization of 53BP1, a pro-NHEJ factor, in cells overexpressing HP1α versus HP1β hints at a probable interplay between HP1 localization and DSB repair protein recruitment. We wondered if this could dictate the differential enrichment of DDR proteins involved in the two main repair DSB pathways, HR and NHEJ, with overexpression of different HP1 isoforms. A different role of HP1β from HP1α in this regard has not been explored in heterochromatin DSBs. We transfected cells with GFP-HP1α or GFP-HP1β, induced damage by laser irradiation in heterochromatin, and performed immunofluorescence (IF) for HR or NHEJ markers such as NBS1, RAD51, phospho-RPA2 (pRPA2), BRCA1, and XRCC4. The cells overexpressing HP1α were largely comparable to untransfected control cells and showed recruitment of early and late HR markers like NBS1, RPA2, RAD51, and BRCA1 (Fig. 2B, C, D, E). Meanwhile, the cells overexpressing HP1β showed impaired enrichment of HR markers but increased recruitment of NHEJ markers like XRCC4 (Fig. 2F) and 53BP1 (Fig. 2A). Conversely, HP1α overexpressing cells showed impaired recruitment of NHEJ marker XRCC4 (Fig. 2F) and 53BP1 (Fig. 2A) at the site of damage. The pro-HR roles of HP1α and pro-NHEJ roles of HP1β were corroborated by performing converse experiments under siRNA-mediated knockdown of these HP1 isoforms. IF for pRPA2 and RAD51 (HR) and XRCC4 (NHEJ) in the background of HP1α or HP1β downregulation mediated by siRNA was performed. HP1β downregulation impaired XRCC4 recruitment compared to scramble control and HP1α siRNA, while HP1α downregulation led to reduced recruitment of pRPA2 and RAD51 compared to scramble control and HP1β siRNA (Supp. Fig. 3A, B). This again indicates the pro-HR role of HP1α and the pro-NHEJ role of HP1β.

**Fig. 2.**
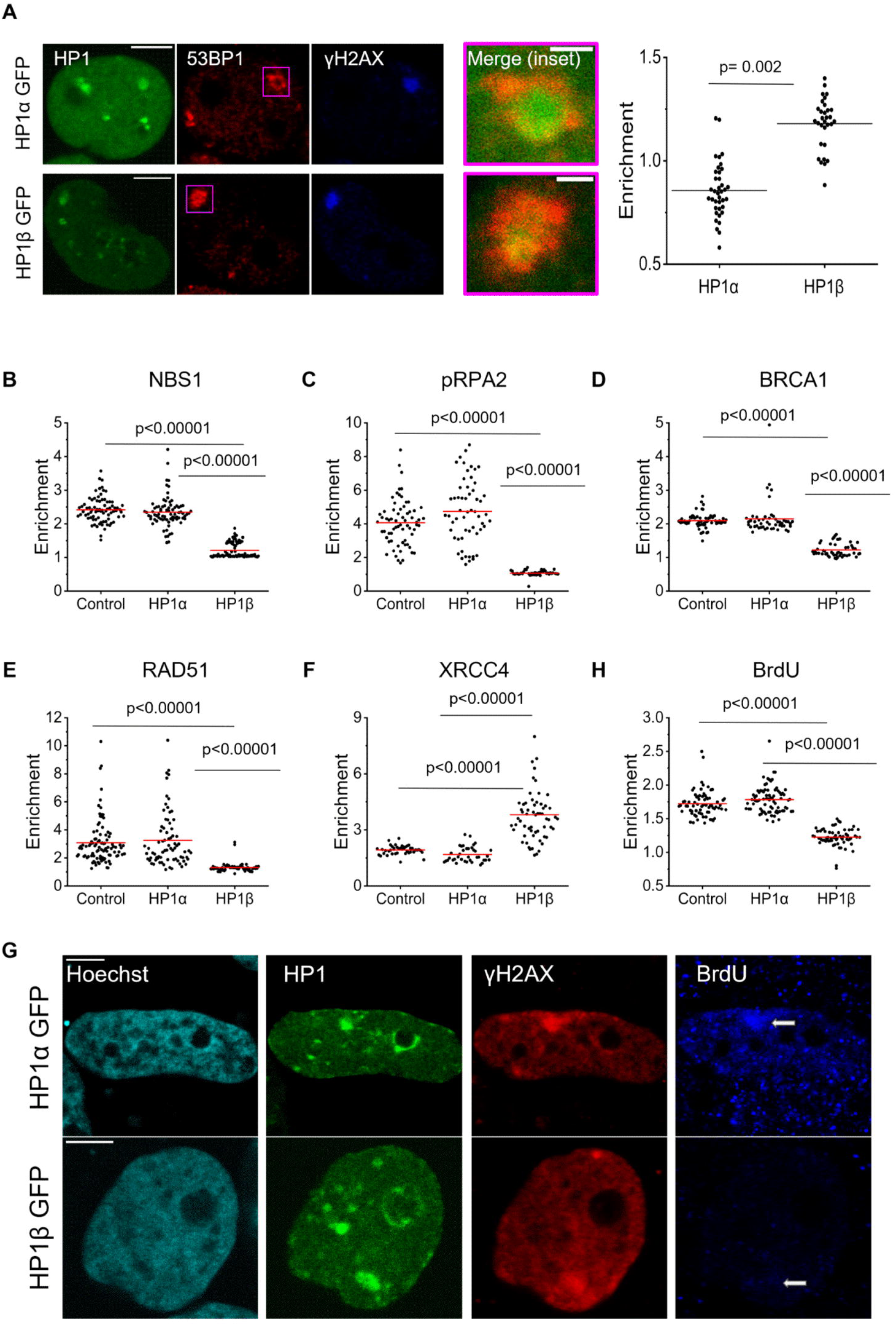
HP1 dynamics can directly influence repair pathway choice at the DSBs in the heterochromatin. (A) Representative images (left) from the immunofluorescence experiment showing 53BP1(red) localization at the site of damage in cells transfected with GFP-tagged HP1α (top) and HP1β (bottom). Localized spot-bleaching was performed within the region marked by the magenta box. The site of damage is marked by γH2AX (blue). The scale bar is 5μm. The merged image shows a zoomed in view of the magenta box. Scale bar for the merge (inset) is 1μm. Quantification of 53BP1 enrichment at the site of damage is shown on the right for HP1α-GFP or HP1β-GFP overexpression. The distribution of cells combined from three immunofluorescence experiments is shown (N=3, n>70 cells, each condition). The line depicts the mean. The p-value is calculated using the Kolmogorov-Smirnov test. Enrichment here is defined as the ratio of intensity at the site of damage to intensity in the ring/annulus around the site of damage (see Supp. Fig 2 for details). Enrichment values < 1 indicate depletion, while enrichment values >1 indicate enrichment. (B-F) Scatter plot showing DDR factor enrichment at the site of damage in control cells and cells transfected with GFP-tagged HP1α or HP1β. The distribution of cells combined from three immunofluorescence experiments is shown (N=3, n>50 cells, each condition). The line depicts the mean. The p-value is calculated using the Kolmogorov-Smirnov test. Enrichment is defined as the ratio of intensity at the site of damage to the intensity in the whole nucleus. Enrichment is shown for (B) NBS1, (C) pRPA2, (D) BRCA1, (E) RAD51, (F) XRCC4 (G) Representative images from immunofluorescence experiment showing BrdU localization (in blue indicated by white arrow) at the site of damage in cells transfected with HP1α (top) and HP1β (bottom). The site of damage is marked by γH2AX (red). The scale bar is 5μm. (H) Scatter plot showing BrdU enrichment at the site of damage in control cells and cells transfected with GFP-tagged HP1α or HP1β. The distribution of cells combined from three immunofluorescence experiments is shown (N=3, n>65 cells, each condition). The line depicts the mean. The p-value is calculated using the Kolmogorov-Smirnov test. Enrichment is defined as the ratio of intensity at the site of damage to the intensity in the whole nucleus.

Beyond the recruitment of HR factors, we confirmed that even the process of HR was affected by these HP1 isoforms. When cells are cultured in the presence of BrdU (an analogue of thymidine), it gets incorporated into the DNA (Latt, 1973). During DDR, when single-stranded DNA is exposed due to resection (specifically during HR), the incorporated BrdU can be detected using an anti-BrdU antibody, which would otherwise remain undetectable in the native condition due to the double-stranded nature of DNA (Cruz-García et al., 2014; Gómez-Cabello et al., 2022; Mukherjee et al., 2015; O’Sullivan et al., 2021; Sartori et al., 2007). Thus, we employed this as a readout for DNA end resection, and consequently, the HR repair pathway. We confirmed that DNA resection occurred at the sites of damage in heterochromatin by performing BrdU immunofluorescence in non-denaturing conditions so that we only detect BrdU in the resected regions. We observed that HP1α overexpressing cells showed BrdU enrichment at the site of damage comparable to control cells, while HP1β overexpressing cells showed significantly less enrichment of BrdU (Fig. 2G, H). This proves that the process of DNA resection and HR is more active at the heterochromatin foci in cells with HP1α overexpression. Our data suggests that HP1α and HP1β promote HR and NHEJ pathways, respectively, for the repair of the DSBs in heterochromatin. Additionally, our data also highlights the difference between HP1α and HP1β in regulating DSB repair and possible spatial segregation of the processes at sites of clustered DSBs, with a core of HP1α-mediated domain conducive to HR surrounded by an HP1β-mediated region of pro-NHEJ factors.

### HP1**α**-mediated phase separation can regulate DSB repair

In the previous experiments, we saw that HP1α can regulate the recruitment of HR proteins and support the error-free HR repair pathway in heterochromatin at the core of clustered DSBs. This led us to wonder if the phase-separating property of HP1α may be essential for setting up such a pro-HR domain. Recently, the ability of proteins to phase separate at the sites of DNA damage to locally enhance the concentrations of repair factors has generated a lot of interest (Altmeyer et al., 2015; Chen et al., 2023; Dall’Agnese et al., 2023; Kilic et al., 2019; Liu et al., 2024; Miné-Hattab et al., 2022; Oshidari et al., 2020; Pessina et al., 2021; Spegg & Altmeyer, 2021; Wang et al., 2023). Independently, HP1α has also been shown to undergo liquid-liquid phase separation (LLPS), which has been implicated in the formation and maintenance of heterochromatin foci (Larson et al., 2017; Strom et al., 2017). To address whether the LLPS ability of HP1α could contribute to the recruitment of HR proteins and the repair pathway choice, we utilized mutants of HP1α, which have been proposed to be incompetent for LLPS (Larson et al., 2017). One such mutant is the S11-S14A mutant (Elathram et al., 2023; Her et al., 2022; Hiragami-Hamada et al., 2011; Sales-Gil & Vagnarelli, 2020) and we chose this for two reasons (Fig. 3A): first, the phosphorylation on the S11-S14 residues is important for LLPS (Her et al., 2022; Larson et al., 2017); second, one of the key differences between HP1α and HP1β is the absence of the four serine residues in the N-Terminal Extension (NTE) region of HP1β, which can regulate many differential functions (Canzio et al., 2014; Hiragami-Hamada et al., 2011; Sales-Gil & Vagnarelli, 2020); NTE phosphorylation of HP1α supports its phase separation. Phosphorylation on S11-S14 residues has been shown to regulate the affinity of HP1α and H3K9me3 histone marks, and long-term overexpression of the S11-S14A mutant, which cannot be phosphorylated, has been shown to negatively affect genome stability (Hiragami-Hamada et al., 2011). We separately overexpressed the GFP-tagged HP1α S11-S14A mutant or wildtype (WT) HP1α in the same backbone to serve as a control. The mutant showed a diffuse localization over nuclei (Fig. 3B). Strikingly, upon laser-induced damage to heterochromatin, the mutant HP1α was readily recruited to the site of damage, as seen using FRAP (Fig. 3C), increasing even beyond initial levels (this is not unexpected as the mutant does not show a special preference for heterochromatin unlike the WT HP1α). To confirm the enrichment of the S11-S14A mutant at the site of damage, we cotransfected the cells with GFP-tagged HP1α S11-S14A and PARP1 chromobody, irradiated a region of interest, and fixed the cells post-damage. We could observe a distinct enrichment of GFP-tagged HP1α S11-S14A mutant at the site of damage (marked by PARP1 enrichment) in these cells (Fig. 3B). However, on investigating the recruitment of HR and NHEJ factors in the background of overexpression of WT or S11-S14A mutants, striking differences emerged. In cells expressing the mutant HP1α, we observed reduced enrichment of several HR factors like pRPA2, RAD51, and BRCA1 (Fig. 3F, G, H) and an increase in the recruitment of NHEJ factors compared to WT HP1α (Fig. 3D). We did not observe any differences in the recruitment of NBS1 (Fig. 3E). Additionally, in an experiment investigating the efficacy of HR as described before, we also saw a reduction in BrdU enrichment on the overexpression of the HP1α S11-S14A mutant (Fig. 3I). This data suggests that the phosphorylation on the NTE could be important, not for HP1α enrichment, but for downstream recruitment of HR factors. We also tested other mutants, like the HP1α I165E mutant, which is defective for polymerization (Brasher et al., 2000). This mutant showed recruitment of DDR factors that was comparable to the control and WT HP1α (Supp. Fig. 4A-F). Thus, the recruitment of HR and NHEJ factors is regulated by the phase separation of HP1α, therefore providing a link between the DDR-centric roles and LLPS of HP1α protein.

**Fig. 3.**
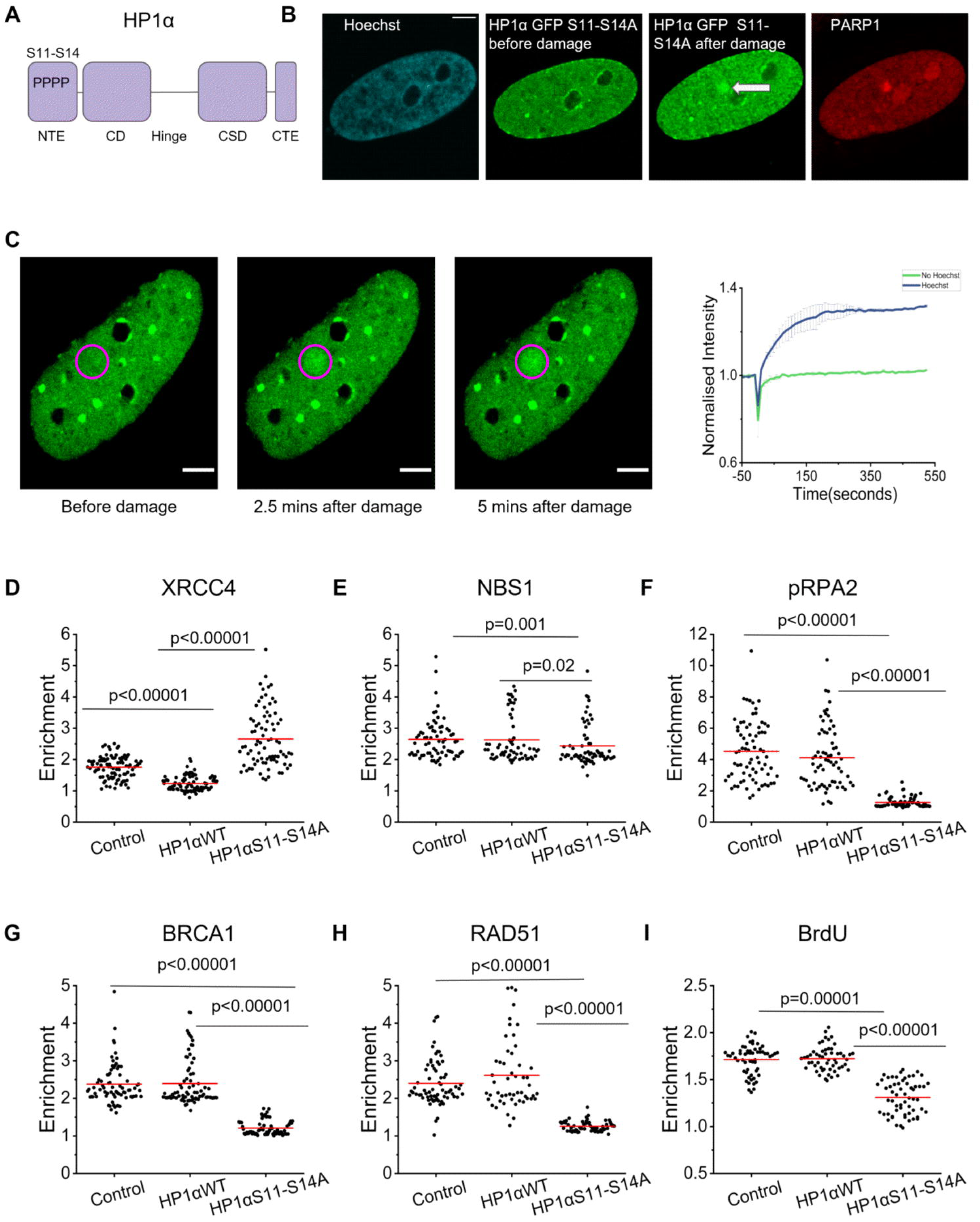
HP1α mediated phase separation can regulate DSB repair. (A) Schematic representing the phosphorylation sites in the N-terminal extension region of HP1α. (B) Images showing GFP-tagged HP1α S11-S14A mutant localization (white arrow) at the site of damage. The site of damage is marked by PARP1 (red). The scale bar is 5μm. DSBs were induced followed by cell fixation and imaging to visualize GFP and PARP1 chromobody. (C) Representative images (left) showing dynamics of GFP-tagged HP1α S11-S14A mutant at different time points of a FRAP experiment with Hoechst sensitization. The magenta circle represents the heterochromatic node that was bleached identified from the Hoechst stain. The scale bar is 5μm. FRAP curve (right) showing the dynamics of GFP-tagged HP1α S11-S14A mutant in undamaged (No Hoechst, green) and damaged cells (Hoechst, blue). The curve depicts the mean and SEM (N=2, n=20 cells, each condition). (D-I) Scatter plot showing DDR factor enrichment at the site of damage in control cells and cells transfected with GFP-tagged HP1α WT or GFP-tagged HP1α S11-S14A mutant. The distribution of cells combined from three immunofluorescence experiments is shown (N=3, n>60 cells, each condition). The line depicts the mean. The p-value is calculated using the Kolmogorov-Smirnov test. Enrichment is defined as the ratio of intensity at the site of damage to the intensity in the whole nucleus. (D) XRCC4, (E) NBS1, (F) pRPA2, (G) BRCA1, (H) RAD51 (I) Scatter plot showing BrdU enrichment at the site of damage in control cells and cells transfected with GFP-tagged HP1α WT or GFP-tagged HP1α S11-S14A mutant. The distribution of cells combined from two immunofluorescence experiments is shown (N=3, n>55 cells, each condition). The line depicts the mean. The p-value is calculated using the Kolmogorov-Smirnov test. Enrichment is defined as the ratio of intensity at the site of damage to the intensity in the whole nucleus.

### HP1**β** dispersal and HP1**α** NTE phosphorylation can directly influence the repair pathway choice

The data suggests that HP1β negatively impacts the recruitment of HR factors (Fig. 2 C, D, E, H), and thus, we expect that its dispersal could be important for their timely enrichment at the sites of damage. Additionally, the potential for phosphorylation on S11-S14 residues of HP1α (and consequent phase separation) seems to positively correlate with the recruitment of HR factors (Fig. 3 F, G, H, I). This led us to investigate if preventing HP1β dispersal after damage could inversely affect the enrichment of HR and NHEJ factors. HP1 proteins undergo diverse PTMs, including phosphorylation, that can regulate the localization and functions of these proteins (Sales-Gil & Vagnarelli, 2020). Phosphorylation of HP1β on T51 residue has been shown to reduce its chromatin binding and allow its dispersal after DNA damage (Ayoub et al., 2008). Additionally, phosphorylation of HP1α on S11-S14 residues is known to tune its affinity to trimethylated histone and directly contribute to its phase separation in heterochromatin (Hiragami-Hamada et al., 2011; Larson et al., 2017). Both of these modifications are performed by the enzyme Casein Kinase II (CKII) (Ayoub et al., 2008; Hiragami-Hamada et al., 2011). We turned to the chemical inhibition of Casein Kinase II (CKII) to inhibit HP1β dispersal post-damage following earlier work (Ayoub et al., 2008). This treatment would also prevent the phosphorylation of the S11-S14 residues in the NTE of HP1α (Nishibuchi et al., 2014) (Fig. 4A). CKII can be inhibited chemically using 4,5,6,7-tetrabromobenzotriazole (TBB) (Ayoub et al., 2008; Hiragami-Hamada et al., 2011; Sales-Gil & Vagnarelli, 2020). We used TBB to inhibit CKII, performed laser-irradiation, and tested the recruitment of HR factor RPA2 and NHEJ factor XRCC4. We expect to see an increase in NHEJ factor enrichment and a reduction in HR factor enrichment if HP1β dispersal and S11-S14 phosphorylation of HP1α are important for such differential recruitment. Compared to control, upon CKII inhibition, the enrichment of pRPA2 was reduced while that of XRCC4 was increased (Fig. 4B, C, D, E). This agrees with the repair factor immunofluorescence data obtained using the overexpression of HP1β and the S11-S14A mutant (Fig. 2 B, C, D, E; Fig. 3 D, E, F, G). This directly implicates HP1β dispersal in regulating the recruitment of NHEJ factor while also corroborating the importance of S11-S14 phosphorylation in affecting HR factor recruitment at DSBs in heterochromatin.

**Fig. 4.**
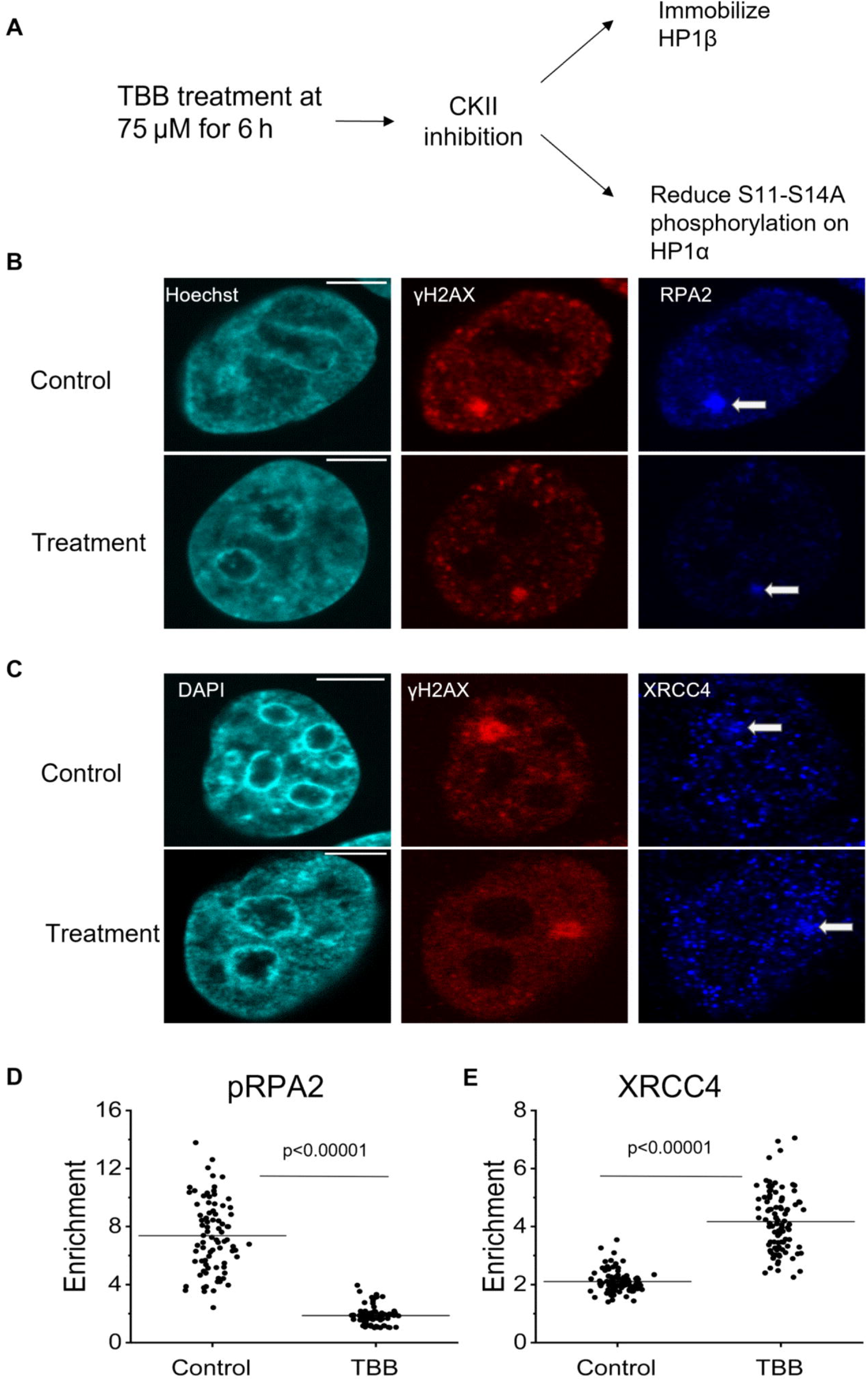
Regulating HP1 dynamics can directly affect the repair pathway choice. (A) Schematic showing the effect of TBB treatment (75μM for 6 hours) on HP1α and HP1β. (B) Representative images from immunofluorescence experiment showing γH2AX (red) and RPA2 (blue) localization at the site of damage in control cells (top) and TBB-treated cells (bottom). Scale bar is 5μm. (C) Representative images from immunofluorescence experiment showing γH2AX (red) and XRCC4 (blue) localization at the site of damage in control cells (top) and TBB-treated cells (bottom). Scale bar is 5μm. (D) Scatter plot showing pRPA2 enrichment at the site of damage in control cells and TBB-treated cells. The distribution of cells combined from three immunofluorescence experiments is shown (N=3, n>100 cells, each condition). The line depicts the mean. The p-value is calculated using the Kolmogorov-Smirnov test. Enrichment is defined as the ratio of intensity at the site of damage to the intensity in the whole nucleus. (E) Scatter plot showing XRCC4 enrichment at the site of damage in control cells and TBB-treated cells. The distribution of cells combined from three immunofluorescence experiments is shown (N=3, n>100 cells, each condition). The line depicts the mean. The p-value is calculated using the Kolmogorov-Smirnov test. Enrichment is defined as the ratio of intensity at the site of damage to the intensity in the whole nucleus.

### The site of damage in heterochromatin represents a phase-separated domain

The data above suggests that the site of damage in heterochromatin is characterized by the accumulation of an LLPS-competent protein, HP1α, that can regulate the repair process. The above experiments using an LLPS-incompetent mutant of HP1α suggest a crucial role for phase separation in driving repair factor recruitment and pathway choice. Phase separation of phosphorylated HP1α has been extensively proven in vitro and using computational models (Elathram et al., 2023; Her et al., 2022; Larson et al., 2017). Beyond puncta formation in cells, other assays are needed to prove phase separation inside cells. Among these are experiments where the dynamics of the protein puncta are tracked within cells as it undergoes liquid-like behaviour, including fission, fusion, and fast recovery in FRAP experiments (Alberti et al., 2019; Titus & Kooijman, 2021). Additionally, cells can be treated with the aliphatic alcohol 1,6-Hexanediol, which has been demonstrated to reduce LLPS by impairing the weak molecular interactions that drive LLPS (Kroschwald et al., 2017); although this can severely affect cellular morphology and machinery (Itoh et al., 2021). Another way to test for LLPS includes tracking the exclusion of an inert free-floating probe, which is kept excluded from the phase-separated puncta due to the absence of specific interactions (Alberti et al., 2019; Larson et al., 2017). HP1 domains are stable and do not undergo dynamic fission or fusion over short time scales (Larson et al., 2017). Hexanediol treatment can affect cell morphology and have other pleiotropic effects, as mentioned above. Thus, to confirm that the site of damage in heterochromatin is indeed a phase-separated domain, we turned to an inert probe exclusion assay performed to confirm LLPS. In this assay, an unrelated and inert probe like EGFP can be excluded from the site where condensates are formed (Fig. 5A). We transfected the cells with a plasmid expressing free-floating EGFP and performed irradiation in heterochromatin to track the EGFP signal over time. There was a steady decrease in the intensity of EGFP at the site of damage, suggesting that phase separation could be operational (Fig. 5B, C). PARP is the early damage sensor implicated in the formation of phase-separated droplets at the site of damage through the formation of chains of PAR polymers that drive phase separation (Altmeyer et al., 2015). To probe the effect of PARP inhibition on EGFP dispersal, we treated the cells with PARPi AZD2461. There was no change in the EGFP exclusion, ruling out PARP as the main mediator of LLPS in this context (Supp. Fig. 5A). Next, we probed the role of HP1α S11-S14 phosphorylation in mediating condensate formation in heterochromatin. To this end, we transfected cells with FLAG-tagged HP1α S11-S14A mutant or FLAG-tagged WT HP1α along with EGFP and performed laser irradiation. The overexpression of the S11-S14A mutant showed EGFP dispersal comparable to the untransfected control, but the overexpression of WT HP1α increased the dispersal of EGFP compared to the untransfected control (Fig. 5C, D). This indicates that the site of damage is a phase-separated condensate, which is mediated by HP1α and its phosphorylation. To specifically tease apart the role of HP1α-mediated LLPS in assisting damage repair and maintaining genomic stability, we overexpressed the S11-S14A mutant and looked at repair defects in cells by Terminal Deoxynucleotidyl Transferase dUTP Nick End Labelling (TUNEL) staining. TUNEL assays can give a direct readout for DNA strand breaks independent of repair protein recruitment. Generally, TUNEL staining is used to label the 3′-OH termini of DNA strand breaks generated during apoptosis-induced DNA fragmentation (Darzynkiewicz et al., 2008). Free 3′-OH is also generated on single-strand or double-strand breaks created during DNA damage (Darzynkiewicz et al., 2008; Figueroa González & Pérez Plasencia, 2017). Thus, the TUNEL assay can be used to measure the strand breaks directly and consequently their disappearance as repair proceeds (Whelan et al., 2020). We found that in cells transfected with WT HP1α, TUNEL intensity went up after damage and came down post 20 hours of recovery comparable to untransfected control, but cells transfected with HP1α S11-S14A mutant showed a repair defect as the TUNEL intensities persisted significantly higher even after 20 hours of damage (Fig. 5E, F). This suggests that the HP1α S11-S14A mutant can hamper the DSB repair process at the site of damage in heterochromatin despite recruiting NHEJ factors.

**Fig. 5.**
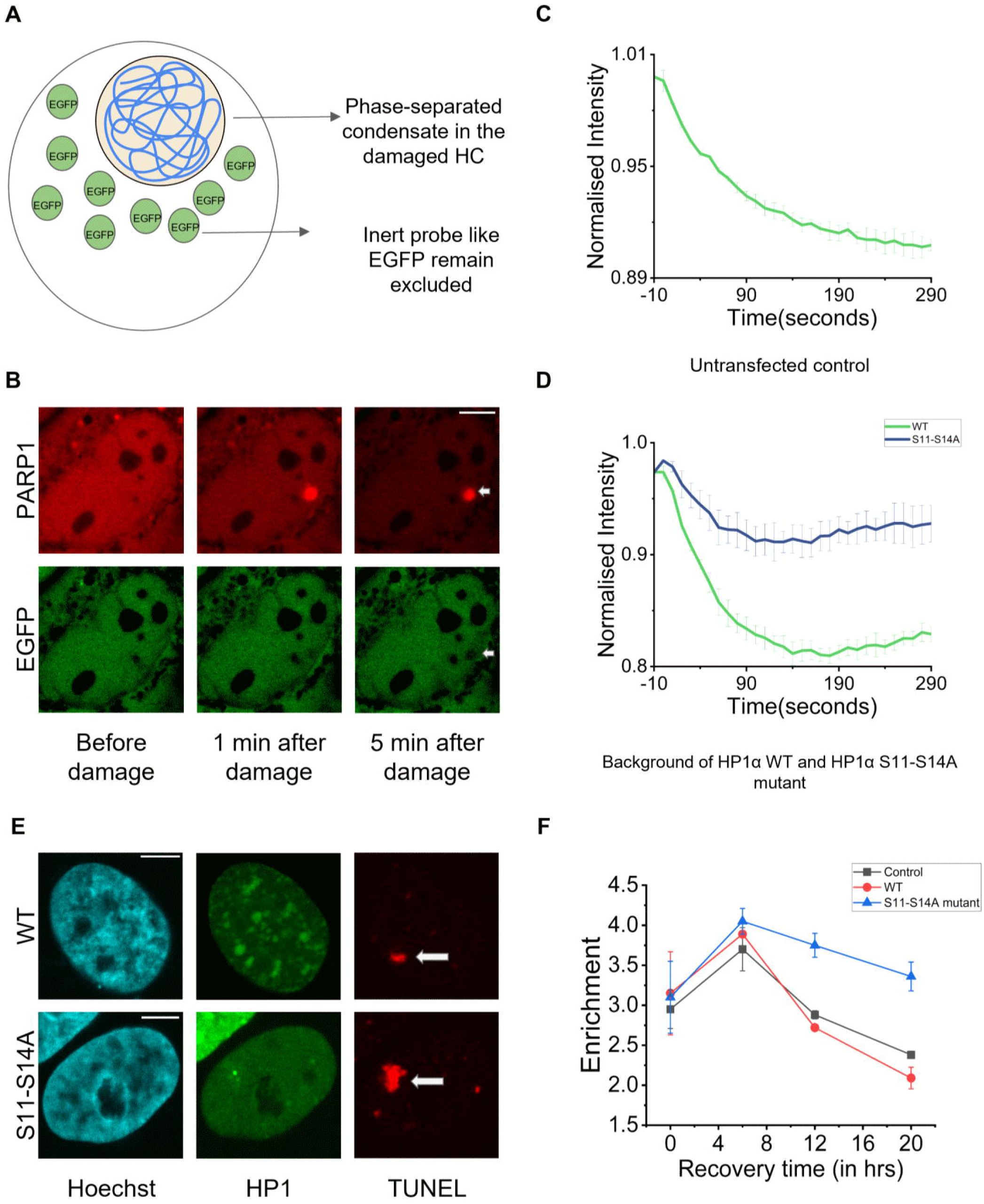
The site of damage in heterochromatin represents a phase-separated domain. (A) Schematic representing the phenomenon of exclusion of free GFP from the phase-separated domains. (B) Images showing the exclusion of EGFP (white arrow) from the site of damage at different times after damage. The site of damage is marked by PARP1(red). The scale bar is 5μm. (C) The exclusion of EGFP from the site of damage is quantified for untreated WT cells. Mean and SEM from two experiments are represented (N=2, n>30 cells, each condition). (D) The exclusion of EGFP from the site of damage is quantified for cells overexpressing HP1α WT (green) and HP1α S11-S14A mutant (blue). Mean and SEM from two experiments are represented (N=2, n>18 cells, each condition). (E) Representative images from the TUNEL assay performed 20 hours after laser-irradiation. Cells with GFP-tagged HP1α WT (top) and GFP-tagged HP1α S11-S14A mutant (bottom) overexpression are shown. Arrow shows the enrichment of TUNEL depicting breaks. Scale bar is 5μm. (F) Repair defect as seen using TUNEL retention detected using TUNEL assay in untransfected control, cells transfected with GFP-tagged WT or S11-S14A mutant. The line graph shows the mean of enrichment at each time point (0, 6, 12, 20 hrs) after damage. Mean and SEM from two experiments are depicted (N=2, n>50 cells, each condition). Enrichment is defined as the ratio of intensity at the site of damage to the intensity in the whole nucleus. The enrichment for WT HP1α and control cells come down at 20 hours but remain higher for S11-S14A mutant.

DSB repair can proceed via two main pathways, namely the HR and NHEJ pathways. HR is a more elaborate and accurate repair pathway, while NHEJ is a more erroneous pathway of repair, resulting in the loss of sequence (Ackerson et al., 2021; Brandsma & van Gent, 2012; Shrivastav et al., 2008). During HR, single-stranded DNA invades a homologous sequence to perform recombination and restore the sequence (X. Li & Heyer, 2008). HR in heterochromatin has specific challenges owing to the repetitive sequences present within the heteroromantic that can undergo unwarranted and illegitimate recombination, leading to genomic instability, and therefore, specific mechanisms exist to safeguard the chromatin (Caridi et al., 2018; Chiolo et al., 2011; Janssen et al., 2018). Based on our data, we put forward another mechanism that can ensure the repair of heterochromatin DSBs while preventing unwanted recombination of damaged sequences with undamaged euchromatin or heterochromatin sequences. We suggest a model in which HP1α enrichment at the site of damage creates an HR-conducive condensate that allows access to HR proteins while HP1β is present at the periphery of this condensate and acts as a limiting element, preventing the spread of HP1α (Keenen et al., 2021). This can limit the region at and around the site of damage where HR can take place. The spatial demarcation created by differential enrichment of HP1α and HP1β can prevent genomic instability by regulating the spatial spread of the HR-conducive region (Fig. 6). Therefore, the establishment of a phase-separated domain at the site of damage in the heterochromatin is of functional importance towards safe repair and the maintenance of genomic integrity.

**Fig. 6.**
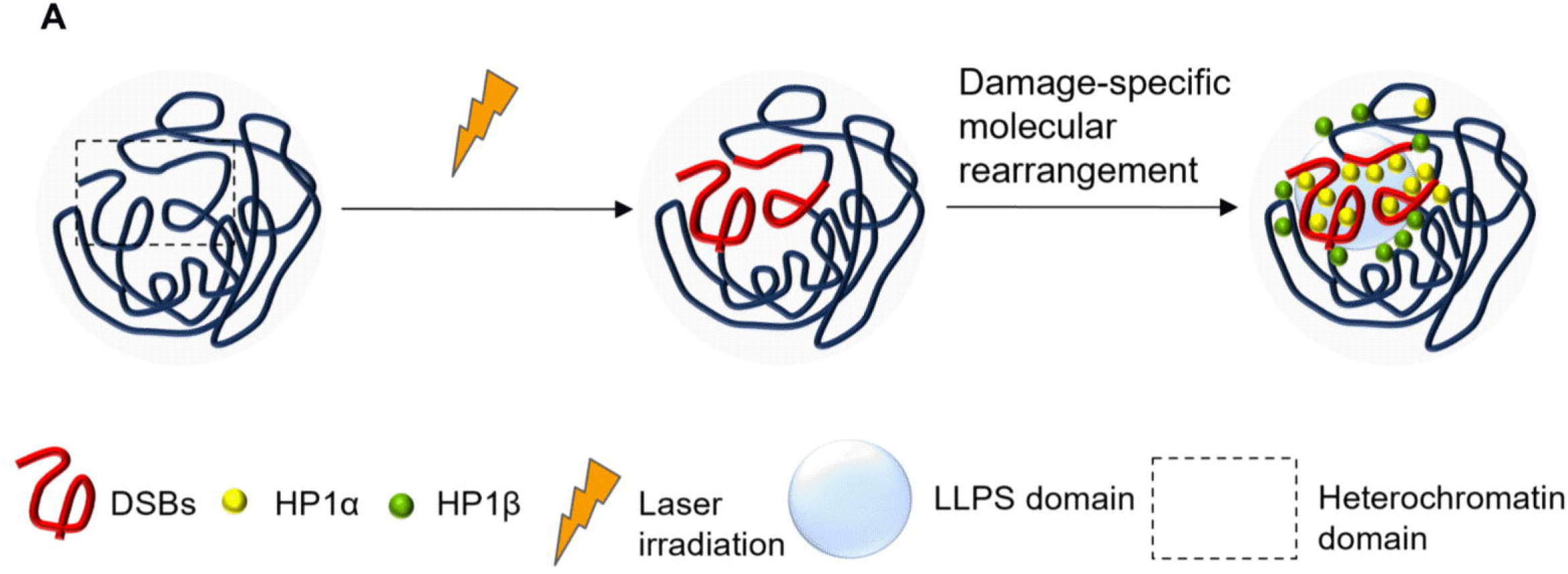
Spatial segregation of HP1 isoforms to allow faithful repair of heterochromatic DSBs. (A) A model depicting the spatial segregation of HP1α and HP1β in response to clustered DSBs in heterochromatin. HP1α mediates LLPS and faithful repair via HR without the risk of ectopic recombination. After induction of DSBs, HP1 localisation changes with HP1α (yellow) enrichment at the site of damage and HP1β (green) enrichment around the site of damage (red). HP1α enrichment could potentially seed the formation and/or enhancement of phase separation (blue circle) at the site of damage. This results in differential propensity for HR and NHEJ at and around the site of damage, restricting HR to the core of damaged heterochromatic node where HP1α-mediated LLPS is most prominent.

## Discussion

The process of DDR occurs in the context of chromatin, and it requires several changes to the chromatin structure to ensure timely repair, followed by restoration of the structure (Dabin et al., 2023; Luijsterburg & van Attikum, 2011; Soria et al., 2012). Based on the condensation status, chromatin can be described as euchromatin or heterochromatin (Bell et al., 2023; Hennig, 1999; Murakami, 2013). The onset of damage on exposure to various sources of damage, as well as the process of repair, have been discovered to be distinct between euchromatin and heterochromatin(Burgess et al., 2014; Goodarzi et al., 2008, 2010; Han et al., 2016; Jakob et al., 2011; Janssen et al., 2016). Questions remain about how the DSBs are detected and repaired in the compact heterochromatin. This is also important because the repetitive nature of chromatin in heterochromatin poses an additional challenge to the repair machinery by being at risk of illegitimate recombination (Janssen et al., 2016). Thus, there is a need to ensure the safe progression of repair pathways like HR and NHEJ without compromising the nature and architecture of heterochromatin sequences (Fortuny et al., 2021). This has been suggested to occur via conserved mechanisms in *Drosophila* and mouse cells (Amaral et al., 2017). We wanted to understand the specifics of DSB repair in heterochromatin and the role of HP1 proteins in such repair. We investigated the roles of the HP1 isoforms that localize to the heterochromatin in mammalian cells. We observed that these architectural proteins that form and maintain the heterochromatin structure are directly involved in heterochromatin DDR. Using laser irradiation to target DSBs in heterochromatin nodes, we investigated the dynamics of the HP1 proteins and discovered an isoform-specific response (Fig. 1B, C, D). This is in agreement with past data but extends them further to reconcile observations in different studies (Ayoub et al., 2008; Baldeyron et al., 2011; Luijsterburg et al., 2009). Here, we pay specific attention to the heterochromatin context and tease apart the isoform-specific roles. Additionally, we find a functional outcome of differential HP1α and HP1β dynamics at the site of damage in terms of the repair pathway choice (Fig. 2B, C). The role of HP1α in DDR and HR has been demonstrated in a different context, where it was shown that p150CAF-1-dependent recruitment of HP1α was essential for the repair of DSBs as it helped in the recruitment of DDR factors such as RAD51 and 53BP1 (Baldeyron et al., 2011). Even though it has been shown HP1 and its isoforms contribute significantly to DNA damage response (Ayoub et al., 2008; Chiolo et al., 2011; Luijsterburg et al., 2009) and HP1α accumulation and its contribution to DSB repair via HR has been established earlier (Baldeyron et al., 2011), this has not been connected to HP1α phase separation. Several studies have shown that phase separation is important for the process of DNA damage repair by assisting in the compartmentalization and recruitment of specific protein complexes (Spegg & Altmeyer, 2021), as in the case of poly(ADP-Ribose) mediated by PARP (Altmeyer et al., 2015), 53BP1 (Altmeyer et al., 2015; Kilic et al., 2019), RAD52 (Oshidari et al., 2020), RAP80 (Qin et al., 2023). Additionally, the role of HP1α phase separation in the formation and maintenance of heterochromatin has been well-explored in the past few years (Elathram et al., 2023; Keenen et al., 2021; Larson et al., 2017; Strom et al., 2017). The role of N-terminal extension (NTE) residues and hinge residues in assisting HP1α phase separation has also become understood using mutants and via simulation studies (Elathram et al., 2023; Keenen et al., 2021; Larson et al., 2017). However, a direct link between HP1α phase separation and DSB repair has not been established thus far. Beyond HP1α, the direct contribution of HP1β in NHEJ has remained elusive as it was shown that knockdown of any HP1 isoforms, including HP1β, did not significantly affect NHEJ efficiency (Alagoz et al., 2015; Lee et al., 2013). While knockdowns may not affect overall efficiency, but still individual HP1 isoforms can finetune the process of HR and NHEJ factor recruitment in specific chromatin contexts. In a different study, no direct impact of HP1α or HP1β on NHEJ could be detected, while HP1γ was implicated, in the context of Cas9-mediated damage in mouse cells with well-defined chromocenters that show repeat clustering (Tsouroula et al., 2016); the differential roles and dynamics of HP1α or HP1β was not described. A subsequent study from the same group indeed showed that heterochromatin DSB response in human cells can be quite different from mouse cells in which the condensed chromocenters can be refractory to HR factors (Mitrentsi et al., 2022). We show a direct role of HP1β in the differential accumulation of pro-NHEJ factor 53BP1 as well as the recruitment of XRCC4 at the site of damage, which is distinct from the pro-HR roles of HP1α in the heterochromatin. To our knowledge, such a differential localization and interaction with DSB proteins has not been shown previously in the context of localized heterochromatin DSB repair. This also supports the fact that DSBs in heterochromatin can recruit HR and NHEJ protein in a positionally stable and spatially partitioned manner.

Thus, HP1α-mediated phase separation has been described in terms of its roles in heterochromatin maintenance, DNA compaction, and transcriptional repression (Her et al., 2022; Keenen et al., 2021; Larson et al., 2017; Strom et al., 2017). How HP1α phase separation impacts the DDR process has not been explored. We investigated HP1α mutants that should potentially be defective for LLPS. We observed that LLPS defects could translate to impaired recruitment of HR proteins (Fig. 3F, G, H). Many DDR proteins have been shown to directly influence or mediate the process of LLPS and regulate downstream repair processes(Altmeyer et al., 2015; Chen et al., 2023; Kilic et al., 2019; Miné-Hattab et al., 2022; Oshidari et al., 2020; Qin et al., 2023; Spegg & Altmeyer, 2021; Wang et al., 2023), and here we connect the hitherto unknown roles of HP1α LLPS to DDR. We directly show the link between HP1α NTE-mediated phase separation (using S11-S14A mutant) and the enrichment of pro-HR factors at the site of damage and its implication in DSB resolution (as shown using TUNEL assay). We speculate that condensate formation is useful for the process of HR by effectively concentrating the HR proteins at the site of damage while preventing the entry of NHEJ proteins into the phase. At the same time, HP1β can regulate the spread of the HP1α seeded phase and promote pro-NHEJ activities if needed. This is possible because past studies have shown that HP1β could dissolve the preformed HP1α droplets (Keenen et al., 2021). Thus, the localization of HP1β would be important to control the spatial localization of HP1α droplets and, in turn, regulate the process of HR at and around the site of damage. The importance of regulating the reach of HR is paramount, especially in the context of heterochromatin. Heterochromatin regions consist of highly repetitive sequences, and the process of HR involves finding a homologous partner. Thus, owing to the repetitive nature of heterochromatin, it can undergo recombination with similar proximate sequences in the genome. This can induce genomic instability and cellular death. In fact, many studies have shown the existence of special systems in place to allow for the faithful repair of heterochromatic DSBs(Caridi et al., 2018; Chiolo et al., 2011; Mitrentsi et al., 2022; Ryu et al., 2015). For instance, in *Drosophila*, the breaks in the heterochromatin migrate outside of the domain to the nuclear periphery to undergo HR repair (Caridi et al., 2018; Chiolo et al., 2011; Ryu et al., 2015). Such relocalization to the periphery has been shown to occur in the case of mouse heterochromatin DSBs (Caridi et al., 2018; Jakob et al., 2011) as well. Strikingly, human heterochromatin DSBs have been shown to be positionally stable and thus, we speculate that HP1α-mediated phase separation could be a way to regulate HR repair in the heterochromatin of human cells. This is of importance, as such protective mechanisms of heterochromatin repair in human cells have remained elusive despite being well understood in drosophila and mouse cells(Caridi et al., 2018; Chiolo et al., 2011; Tsouroula et al., 2016).

Another question of relevance is why the heterochromatin sequences choose to undergo HR, given the risk it involves. The answer to that could lie in the nature of the DSBs that are being induced. The choice of repair pathway is dictated by many factors, and one such factor is the complexity of the break. It has been shown that complex breaks requiring end processing tend to undergo HR not NHEJ (Shibata et al., 2011, 2014).

We propose that the clustered DBSs induced in the human heterochromatin by laser irradiation could necessitate using HR pathway and LLPS acts to prevent illegitimate recombination and genomic instability. In this context, it is important to point out that in highly metastatic cancers, a downregulation of HP1α has been observed (Kirschmann et al., 2000; Norwood et al., 2004). More studies are required to link the LLPS of HP1α to its direct role in the maintenance of genomic stability, especially in the context of DNA damage and associated pathologies.

## Materials and methods

### Cell culture

U2OS cells were obtained from ATCC (via HiMedia Laboratories). U2OS cells were grown in McCoy 5A media (Sigma, M4892) supplemented with 10% FBS (Gibco, 16000-044) and 1% PenStrep-glutamine (Gibco, 10378-016). HeLa cells were obtained from NCCS. HeLa cells were grown in DMEM/F12 (Gibco, 12400-024) supplemented with 10% FBS (Gibco, 16000-044) and 1% PenStrep-glutamine (Gibco, 10378-016). Cells were grown in T-25 flasks (Tarsons, 950040) and incubated in a CO_2_ incubator (Eppendorf Galaxy 170S) set to 37°C and 5% CO_2_. For experiments and imaging, the cells were plated in a glass-bottom dish (Genetix, 200350) at least 24 hours before the experiments. For live cell experiments, the cells were imaged directly in a supplemented FluoroBrite medium (Gibco, A1896701). The plates were placed in the live-cell chamber and maintained at 37°C. Transfection was performed using the X-tremeGENE™ HP DNA Transfection Reagent (Roche, 6366236001) following the manufacturer’s protocol.

### Immunofluorescence (IF)

The cells were grown as described previously. Following damage, cells were allowed to recover (if mentioned) and afterward fixed using 4% PFA (paraformaldehyde, Sigma P6148) in 1X PBS (phosphate-buffered saline, Sigma P5493) for 10 minutes. After 10 minutes, PFA was removed, and cells were washed twice with 1X PBS. Permeabilization was performed using 0.3% Triton X-100 (Sigma, T8787) in 1X PBS for 10 minutes. Triton was removed and cells were washed twice with 1X PBS. After permeabilization, the cells were blocked with 5% BSA (bovine serum albumin, Himedia TC545) in 1X PBS (Blocking solution) for an hour. This was followed by the treatment with primary antibodies (with dilution as mentioned in the table) in the blocking solution. Incubation with primary antibodies was done overnight at 4°C. On the next day, the primary antibodies were removed from the plate and cells were washed thrice with 1X PBS. After this, the cells were incubated with the secondary antibodies in the blocking solution for 2 hours at RT. Secondary antibodies were washed out and cells were washed thrice with 1X PBS. Finally, the cells were stained with DAPI (1ug/ml in 1X PBS) for 10 minutes. The cells were imaged in 1X PBS unless mentioned otherwise.

### Inhibitors and treatments

The inhibitors and their concentrations are as follows: PARP inhibitor AZD2461 was used at 60μM for 24 hours. CKII inhibitor TBB (Sigma, 218697) was used at 75μM for 6 hours. BrdU (Sigma b5002) was used at 5μM for 24 hours.

### EGFP exclusion

Free probe exclusion was used to assess LLPS condensates. EGFP was overexpressed followed by irradiation of a heterochromatic node with 405nm laser in Hoechst sensitized cells. EGFP intensity was tracked over time. Intensity at the site of damage normalized by the intensity in the nuclei is plotted (like FRAP curves). In the case of the WT and S11-S14A mutant experiment, FLAG-tagged HP1α WT or S11-S14A mutant was overexpressed along with EGFP.

### TUNEL assay

The assay was performed using the In Situ Cell Death Detection Kit, TMR red (Roche, Cat. No. 12156792910). The kit helps label and quantify the DNA strand breaks at single-cell level. The principle is based on labelling the free end of SSB or DSB using the enzyme Terminal deoxynucleotidyl transferase followed by detection using fluorescent imaging. The intensity was quantified on a cell-by-cell basis at each time point after damage to quantify the persistence/repair of damage.

### Confocal microscopy and micro-irradiation

Imaging was performed on a confocal microscope unless mentioned otherwise. Induction of DSBs in Hoechst-sensitized cells with a 405 nm has been described before (Dinant et al., 2007; Kesavan et al., 2020). Fluorescence images were acquired using a 60× oil objective (PlanApo N 60× Oil, numerical aperture = 1.42, Olympus) mounted on an Olympus IX83 inverted microscope equipped with a scanning laser confocal head (Olympus FV3000). Cells grown in 35-mm glass-bottomed dishes were sensitized with 1µg/ml Hoechst for 10 minutes in media. After Hoechst sensitization, cells were incubated in a supplemented FluoroBrite medium before imaging. The plate was positioned on the confocal microscope, point ROI was drawn in the region of interest and DSBs were induced with the 405-nm laser (as described in Dinant et al., 2007 (Dinant et al., 2007)).

### FRAP analysis

FRAP analysis was performed according to Phair and Misteli, 2000 (Phair & Misteli, 2000). Briefly, the intensity at the site of bleach was determined using a circular ROI at different time points. The intensity from 5 images before bleaching was determined and the mean was calculated (referred to as Io, the prebleach intensity). The time-dependent intensities at the site after the bleach were designated as It. Similarly, using a free-hand ROI and image registration, the intensity of the whole nucleus in the 5 images before bleaching was determined and the mean was calculated (referred to as No). The intensities in the nuclei after the bleach were designated as Nt. The factor No/Nt corrects for loss of overall intensity during the bleach pulse and any subsequent photobleaching in the time-course of imaging ^90^. The intensity as the site normalized to the intensity in the nuclei was calculated at each time point given by

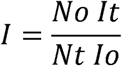

### Wide-field microscopy

For wide-field imaging, the plates were imaged on an Olympus IX83 inverted widefield fluorescence microscope with a Retiga 6000 CCD monochrome camera (QImaging). The images were acquired using a 60X, 1.42 N.A. oil immersion objective or a 100X, 1.4 N.A. oil immersion objective (unless mentioned otherwise).

### Image analysis

The multichannel images were split and renamed using Fiji. For representative images were contrast adjusted in Fiji and shown. Intensity analysis was done either in Fiji or using an in-house MATLAB (Mathworks) script on a cell-by-cell basis.

### Statistical analysis

The graphs were plotted using Python 3, MATLAB, and Origin Pro. To perform a student’s t-test for statistical significance, GraphPad (online) was used. To compare means, quantification was done using data from at least three independent replicates unless mentioned otherwise. The bar graphs represent the mean and standard error of the mean (SEM) unless mentioned otherwise. For single-cell distributions, a non-parametric test (Kolmogorov-Smirnov’s test) was employed as it does not make assumptions regarding the normality of the underlying data. Single-cell distributions may not follow a normal distribution and a non-parametric test is more suited for these cases. p values below 0.05 were considered statistically significant.

### Antibodies and plasmids

The dilutions of all primary and secondary antibodies used for the IF are given in Table S1. The following plasmids were used in this study: GFP-tagged HP1α and HP1β were obtained from Addgene (Addgene #17652 and #17651). GFP-tagged and FLAG-tagged HP1α WT and S11-S14A (Hiragami-Hamada et al., 2011) mutant were received as a gift from Jun-ichi Nakayama. GFP-tagged HP1αWT, HP1αI165E mutant (Shibata et al., 2011), and HP1αW174A mutant were received as a gift from Lori Wallrath. pcDNA3-EGFP was obtained from Addgene (Addgene #13031).

---

**Table S1.**

| Antibody | Dilution | Catalog number |
| --- | --- | --- |
| Mouse γH2AX | 1:1000 | Merck-JBW301 |
| Mouse HP1α | 1:800 | Merck-05-689 |
| Rabbit HP1α | 1:200 | CST-2616S |
| Rabbit HP1β | 1:800 | CST-8676S |
| Rabbit 53BP1 | 1:500 | Abcam-ab175933 |
| Rabbit RAD51 | 1:500 | Abcam-ab221796 |
| Rabbit XRCC4 | 1:500 | Abcam-ab97351 |
| Rabbit PAR | 1:500 | Biotechnie-4335-MC-100 |
| Rabbit pRPA2 | 1:500 | CST-54762 |
| Rabbit NBS1 | 1:100 | CST-14956 |
| Rat BrdU | 1:500 | Abcam-ab6326 |
| Mouse BRCA1 | 1:500 | Merck-SAB2702136 |
| Mouse secondary 488 | 1:1000 | Invitrogen-A11029 |
| Mouse secondary 594 | 1:1000 | Invitrogen-A11032 |
| Mouse secondary 647 | 1:1000 | Invitrogen-A21236 |
| Rabbit secondary 488 | 1:1000 | Invitrogen-A11034 |
| Rabbit secondary 594 | 1:1000 | Invitrogen-A11037 |
| Rabbit secondary 647 | 1:1000 | Invitrogen-21245 |
| Rat secondary 647 | 1:1000 | Invitrogen-A21247 |

## Funding and Acknowledgements

This project was supported by intramural funds at TIFR Hyderabad from the Department of Atomic Energy, Government of India (Project Identification No. RTI 4007). GFP-tagged HP1α and HP1β were a gift from Tom Misteli (Addgene #17652 and #17651). GFP-tagged and FLAG-tagged HP1α WT and S11-S14A mutant ^11^ were a gift from Jun-ichi Nakayama. GFP-tagged HP1αWT and HP1αI165E mutant ^84^ were a gift from Lori Wallrath. pcDNA3-EGFP was a gift from Doug Golenbock (Addgene #13031).

## Declaration of Interests

The authors declare no competing interests.

## Notes

### Competing Interest Statement

The authors have declared no competing interest.

