## Supplementary Information for "HP1α-driven Phase Separation and Repair Pathway Choice in Response to Heterochromatin Damage"

**Supp. Fig. 1**

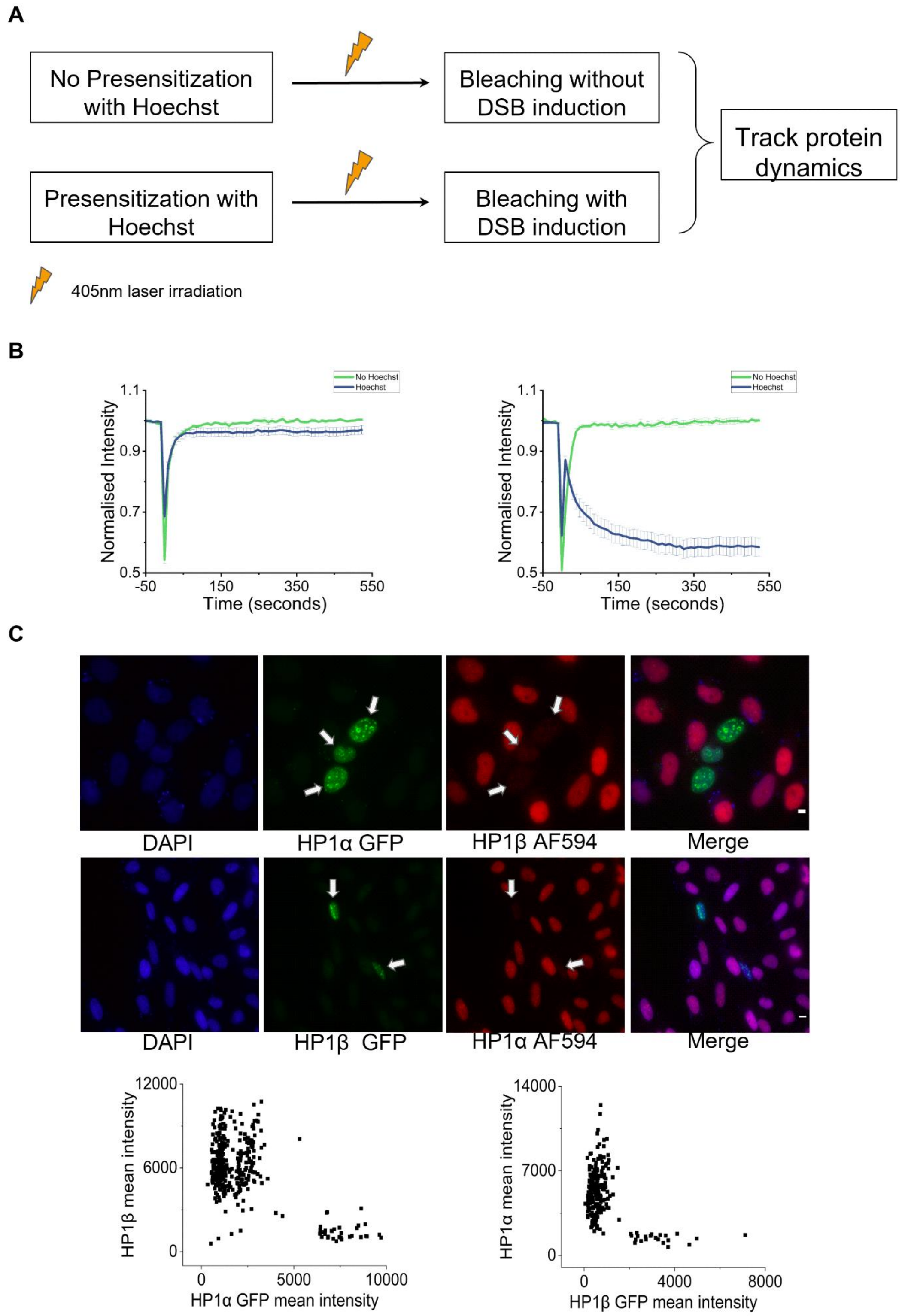
A
No Presensitization with Hoechst
Bleaching without DSB induction
Presensitization with Hoechst
Bleaching with DSB induction
Track protein dynamics
405nm laser irradiation
B
No Hoechst
Hoechst
Normalised Intensity
1.1
0.9
0.7
0.5
-50
150
350
550
Time (seconds)
C
DAPI
HP1α GFP
HP1β AF594
Merge
DAPI
HP1β GFP
HP1α AF594
Merge
HP1β mean intensity
12000
6000
0
5000
10000
HP1α GFP mean intensity
HP1α mean intensity
14000
7000
0
4000
8000
HP1β GFP mean intensity

A. Schematic depicting the process of Hoechst sensitization and laser irradiation adopted to inflict DSBs and study protein dynamics via FRAP simultaneously. DBSs are induced only when Hoechst sensitization is performed.

B. FRAP curves from HeLa cells show the dynamics of HP1α and HP1β in undamaged control (green) and damaged cells (blue). The curve depicts the mean and standard error of the mean (SEM) from one experiment (N=1, n>10 cells).

C. Representative images from the immunofluorescence experiment imaged on a widefield microscope are shown. It depicts the cells (white arrows) with GFP tagged HP1α overexpression showing downregulation of HP1β (top) and vice versa (bottom). The scale bar is 5µm. The mean intensity of HP1 GFP has been plotted against the mean intensity of immunostained HP1 to quantify the correlation between the two (scatter plot). Note that in the images and the quantification how the cells expressing high levels of GFP-tagged HP1α show low levels of HP1β and vice versa.

**Supp. Fig. 2**

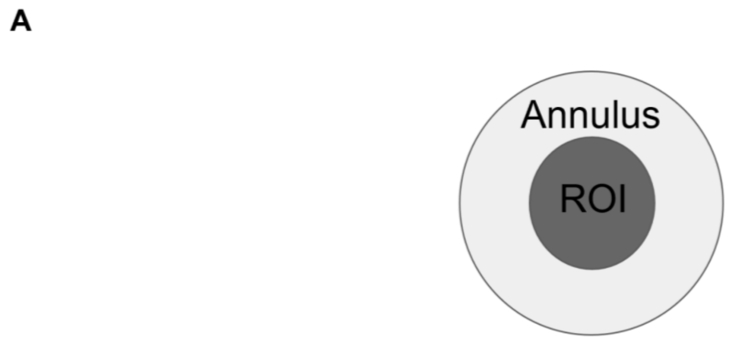


A. Illustration depicting the ROI and annulus used to analyze the data pertaining to FIG. 2A. The ROI was segmented using a damage marker (γH2AX or PARP1), and the ROI was used to find the mean intensity at the site of damage. The ROI was then dilated to create a bigger ROI from which the smaller ROI was removed resulting in the annulus around the site of damage. The mean intensity in the annulus was also calculated. The ratio of mean intensity in the ROI to mean intensity in the annulus was calculated and plotted as enrichment in cases where a ring-like arrangement was apparent.

**Supp. Fig. 3**

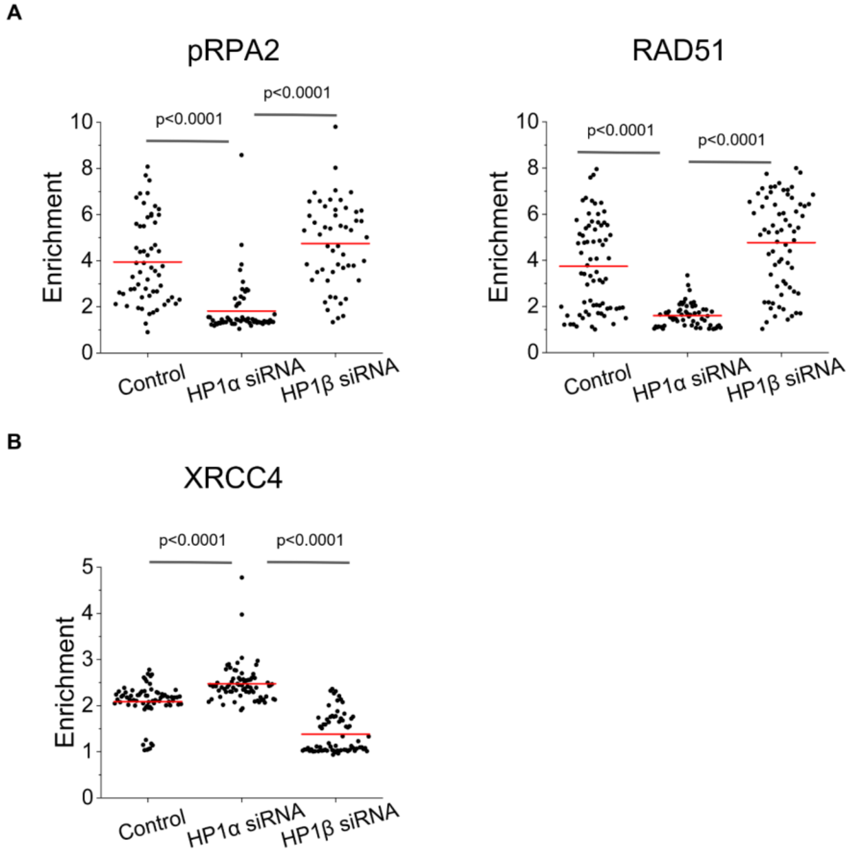


(A-B) Scatter plot showing DDR factor enrichment at the site of damage in cells transfected with scramble siRNA, HP1α siRNA, or HP1β siRNA. The distribution of cells combined from two immunofluorescence experiments is shown (N=2, n>50 cells, each condition). The line depicts the mean. The p-value is calculated using the Kolmogorov-Smirnov test. Enrichment is defined as the ratio of intensity at the site of damage to the intensity in the whole nucleus. (A) pRPA2 and RAD51, (B) XRCC4

**Supp. Fig. 4**

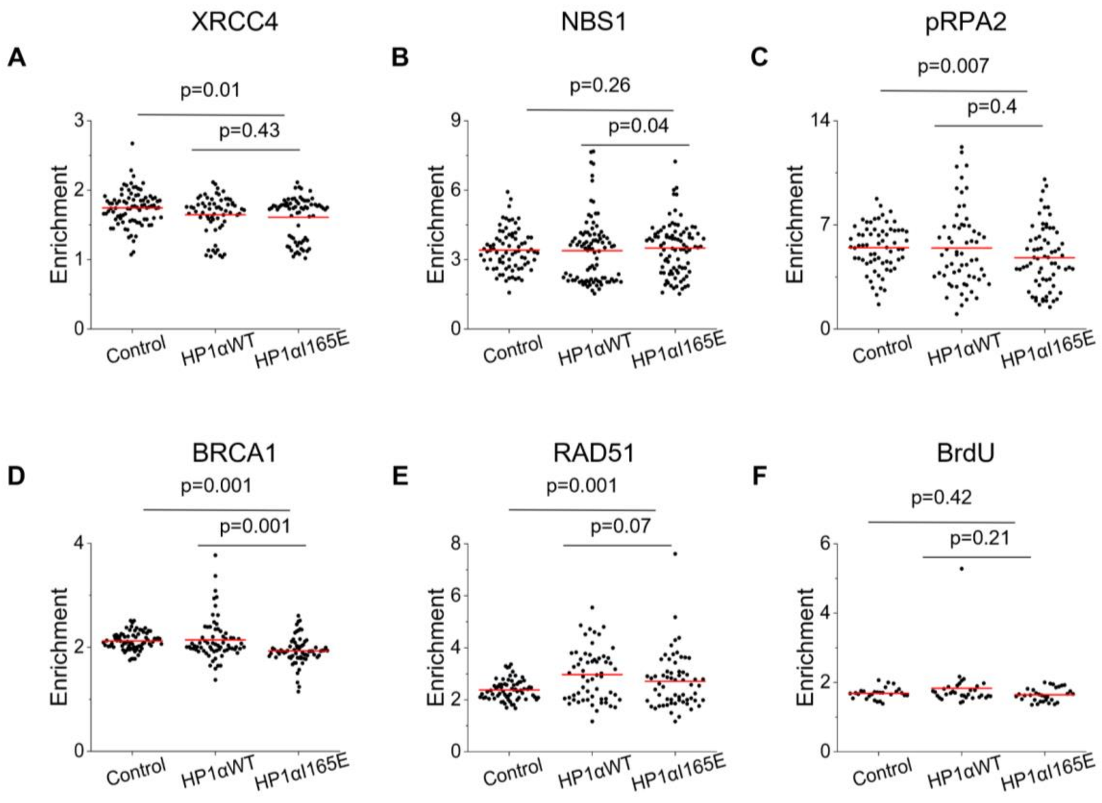


(A-E) Scatter plot showing DDR factor enrichment at the site of damage in control cells and cells overexpressing HP1αWT and HP1αI165E mutant. The distribution of cells combined from three immunofluorescence experiments is shown (N=2, n>60 cells, each condition). The line depicts the mean. The p-value is calculated using the Kolmogorov-Smirnov test. Enrichment is defined as the ratio of intensity at the site of damage to the intensity in the whole nucleus. (A)XRCC4, (B)NBS1, (C) pRPA2, (D) BRCA1, (E) RAD51

(F) Scatter plot showing BrdU enrichment at the site of damage in control cells and cells overexpressing HP1αWT and HP1αI165E mutant. The distribution of cells combined from two immunofluorescence experiments is shown (N=2, n>30 cells, each condition). The line depicts the mean. The p-value is calculated using the Kolmogorov-Smirnov test. Enrichment is defined as the ratio of intensity at the site of damage to the intensity in the whole nucleus.

**Supp. Fig. 5**

A

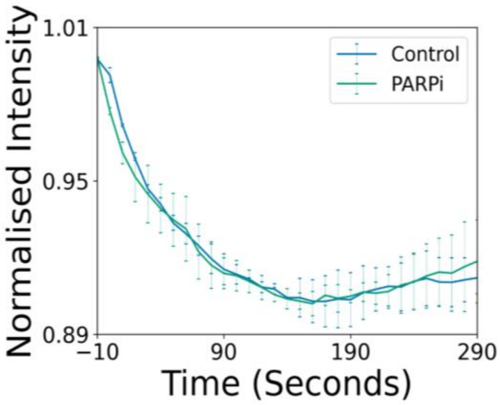


A. The exclusion of EGFP from the site of damage is quantified for control cells (blue) and cells treated with 60uM AZD2461 for 24 hours (green). Mean and SEM from two experiments are represented (N=2, n>20 cells, each condition).
